# Single-cell carbon storage dynamics drive conditional fitness in microbes

**DOI:** 10.64898/2026.04.08.717106

**Authors:** Jiaqi Huang, Ruoshi Yuan, Yihao Ma, Haoming Ma, Adam P. Arkin

## Abstract

Microbes frequently encounter fluctuating environments, requiring dynamic energy management strategies for survival. While carbon storage polymers like polyhydroxybutyrate (PHB) are ubiquitous across bacterial taxa, their precise ecological advantage remains poorly understood.^1^ Here we show that carbon storage drives conditional fitness during environmental transitions. Using a high-throughput single-cell microfluidic platform, we tracked tens of thousands of *Cupriavidus necator* cells under precisely controlled carbon and nitrogen fluctuations. We found that PHB provides no advantage under nutrient abundance but becomes decisive at starvation boundaries: during carbon starvation, it enables ∼30% more progeny before arrest; during recovery from nitrogen starvation, it shortens lag and accelerates regrowth. Strikingly, at the single-cell level, PHB granules are inherited in an asymmetric, all-or-nothing fashion, concentrating resources into specific lineages to overcome the discrete energetic threshold required for cell division. Despite this single-cell variance, at the population level, PHB fractions robustly return to a common setpoint after nutrient shifts—a homeostatic behavior consistent with integral feedback control. These findings reveal that while PHB does not increase the basal exponential growth rate, it confers a distinct fitness advantage by prolonging the proliferative phase during nutrient depletion and facilitating successful recovery from starvation, explaining the evolutionary persistence of carbon storage in environments with pulsed resource availability.

---

Most carbon in the biosphere ultimately passes through microbes, with a fraction retained as storage compounds intracellularly rather than immediately respired to the atmosphere^2,3^. In soils, microbial carbon storage can constitute ∼16–54% of total microbial carbon^4^, buffering cells against nutritional limitation^5,6^, feast–famine cycles^7,8^, and other stresses^9^. Microbes deploy multiple storage strategies matched to limiting elements: for carbon and energy, trehalose^2^, glycogen^10^ , triacylglycerides (TAG)^2^, and polyhydroxyalkanoates (PHAs) such as polyhydroxybutyrate (PHB)^1^; for phosphorus, polyphosphate^11^; for nitrogen, polyamines^12^,and cyanophycin^13^. Importantly, dedicated insoluble storage polymers can reach very high concentrations—PHB up to ∼50–90% of cell dry weight—without increasing osmotic pressure or risking lysis, unlike soluble central-metabolite pools^14^.

PHB is widespread across bacterial clades and accumulates as phase-separated granules with a protein coat. Beyond serving as a carbon/energy reserve, PHB metabolism has been linked to redox homeostasis^15,16^, stress resistance including osmotic buffering^17^, freeze–thaw^18^resilience, and metabolic flexibility^15^. *Cupriavidus necator* is both a prevalent soil bacterium and a model PHB producer^19,20^.

Much of the PHB literature has been shaped by biotechnological goals—optimizing renewable bioplastic yields—yielding detailed knowledge of the granule biogenesis pathway^19,20^and high-yield strategies in fed-batch or two-stage reactors, typically via nitrogen limitation ^2,3,21,22^. These industrial processes leverage a ’High PHB’ metabolic mode induced by an abundance of carbon and high C:N ratio, channeling the surplus acetyl-CoA pool toward PHB accumulation^22–24^. Yet these conditions differ fundamentally from the heterogeneous soils^25,26^ and freshwater-associated habitats where *C. necator* is typically found. These natural habitats are typically oligotrophic, characterized by dilute, chemically diverse nutrients and rapidly turning-over substrate pools^27–29^. A smaller pool of low-molecular-weight substrates—e.g. Sugars, organic acids,^27^ and free amino acids^29^—turn over rapidly^28,29^. Consequently, observations from resource-rich, steady cultivations likely fail to capture the dynamic survival strategies microbes deploy in nature.

Furthermore, most prior work relies on end-point measurements under constant or slowly changing conditions, missing dynamic strategies used to cope with capricious supply. Second, bulk measurements average over single-cell heterogeneity that can be crucial for population persistence^21,22^. Recent single-cell physiology underscores this: “steady state” is best defined by time-invariant single-cell distributions rather than bulk OD^30^, and cells subjected to repeated nutrient fluctuations adopt physiologies that differ from single-shift expectations^31^. Moreover, resource-allocation theory predicts that during transitions, protein synthesis and sector allocation can be suboptimal—a consequence of strategies tuned in response to fluctuating environments rather than instantaneous optimization^32,33^.

To address these gaps, we used a high-throughput microfluidic “mother machine”^34,35^ to trap tens of thousands of *C. necator* cells and track them for days under precisely controlled carbon (C) and nitrogen (N) supply regimes. This design lets us (i) monitor PHB granule accumulation and inheritance across many generations, (ii) separate transient from steady-state responses to nutrient up- and down-shifts within replete conditions, and (iii) directly compare wild-type and

PHB-deficient mutants under matched C and N regimes, as well as very low C and N concentrations. We show that C. necator maintains PHB granules and passes them to daughter cells in an all-or-nothing fashion, generating storage-rich and storage-poor lineages without detectable growth cost—observations at the single-cell that would be obscured in bulk, end-point assays. Under step changes within replete conditions, PHB fractions return to a common setpoint. In control theory, perfect adaptation refers to the return of a regulated variable to its setpoint despite sustained perturbation^36^. Exhibiting exactly this phenomenology, the system’s dynamic is reminiscent of integral-feedback control, and aligns with resource-allocation frameworks in which proteome and precursor limitations impose trade-offs between response dynamics and instantaneous growth in fluctuating environments^32,36^. Across starvation thresholds, PHB shifts from neutral to decisive: it yields extra progeny during transition into stationary phase after carbon depletion and shortens lag while emerging from stationary phase after nitrogen starvation. Together, these time-resolved single-cell dynamics reveal a storage program that is effectively neutral in constant abundance yet strongly beneficial at starvation boundaries, suggesting that PHB-mediated diversification and control confer a fitness advantage in environments with pulsed resource availability.

## Platform & Validation

To analyze single-cell dynamics of *C. necator* growth and PHB production, we modified the “mother machine” microfluidic device^34^ to provide precisely maintained and rapidly switchable growth environments across a wide range of nutrient concentrations. Several problems had to be solved for this platform to deliver robust biological measurements: (i) nutrients must diffuse efficiently into the narrow trenches where cells reside, (ii) long-term experiments must avoid inlet clogging and spatial gradients, (iii) PHB granules must be imaged without perturbing their synthesis, and (iv) the system must support high throughput and automated analysis to capture rare phenotypes.

### Device design and engineering solutions

Our optimized chip features trenches one cell width and height, spaced at 4 µm intervals, with a total of 124,500 trenches per chip (Fig. 1A). To improve nutrient access, we fabricated trenches with a trapezoidal cross-section, increasing effective diffusion area by ∼33% compared to square cross-section profiles. To prevent clogging of the main channel during long experiments, we introduced a three-lane inlet–outlet design: cells are only loaded into the distal lane, while the two upstream lanes remain cell-free, ensuring unobstructed flow (Fig. 1E–G). Trench dimensions were verified using atomic force microscopy (AFM), and finite-difference modeling predicted >95% equilibration of nitrogen within ∼1 second at standard flow rates. These design features allow abrupt, reproducible nutrient shifts at timescales much shorter than the cell cycle. Nutrient shifts through medium switches typically finish in less than a minute.

**Figure. 1.**
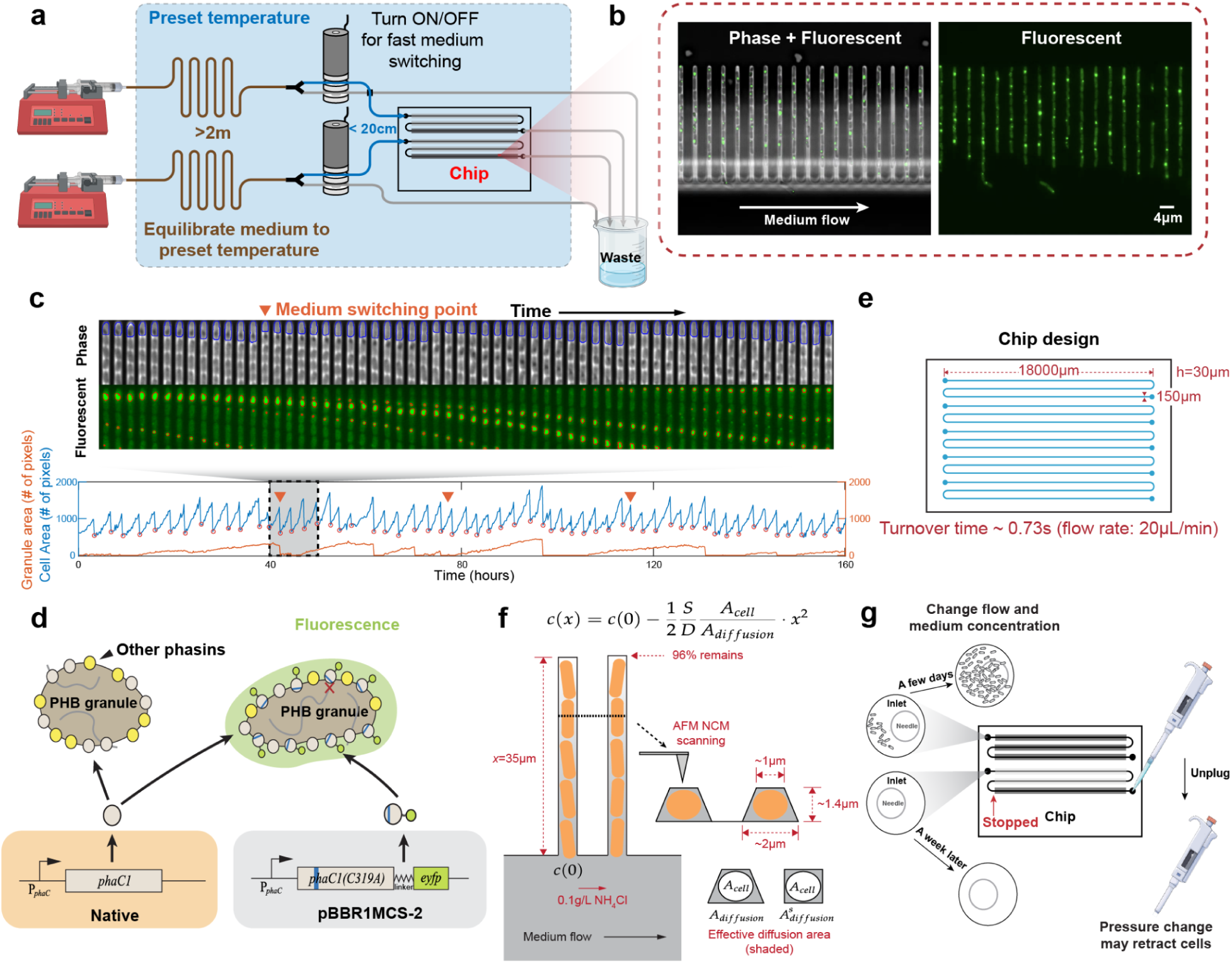
Overview of high-throughput and long-term visualization and quantification of polyhydroxyalkanoates (PHA) in single living cells under perturbation of rapidly switchable environmental conditions. **a.** Microfluidic setup for fast medium switching: Syringe pumps (left) continuously supply medium through a temperature-equilibrated, long tubing (>2 meters) to the chip. A Y-junction with an integrated pinch valve directs flow either to the chip (pinch valve OFF) or to waste (pinch valve ON). **b.** We applied an established single-cell imaging microfluidic design known as the mother machine, which allows cells to grow within single-cell width trenches while a growth medium is pumped through a much larger orthogonal feeding channel. **c.** The growth trajectory of a single cell from a week-long experiment with three medium switches. The upper panel displays the kymograph with 60 frames of the top of its trench in both phase contrast and fluorescent channels. We employed a deep learning-based tool (DELTA) to segment cells from phase contrast images, with identified cell boundaries indicated in blue. **d.** A single amino-acid substitution (C319A) of PHB synthase PhaC1 fused with eYFP is employed to visualize PHB granules. This mutant PhaC1 is unable to synthesize PHB but retains the ability to bind to PHB granules, thus enabling their detection. **e.** Highlighted microfluidic chip design with dimensions from authentic GDSII file for photomask fabrication. The corresponding medium turnover time is about 0.73 seconds under the flow rate of 20μL/min **f.** Atomic force microscopy measurements of the single-cell trenches’ dimensions, revealing a trapezoidal cross-sectional shape. The microscope works in non-contact mode (NCM) to avoid potential damage to trenches. The effective diffusion area, calculated as this trapezoidal area minus the cross-sectional area of a cell, is critical in our model of diffusion transport. Our model suggests a quadratic decrease in nutrient concentration along the trench. **g.** In our exploration of ultra-low nitrogen conditions, we have strategically designed a microfluidic chip featuring a serpentine three-lane structure. By loading cells (condensed culture) into only the last one or two lanes, we avoid condensed cell culture reaching and residing at the chip’s inlet, thereby avoiding nutrient depletion and potential inlet blockage.

### Granule imaging and lineage tracking

To visualize PHB granules, we used an EYFP-tagged, catalytically inactive PhaC1 (dPhaC1-EYFP) expressed from its native promoter on a low-copy plasmid^37^. This strategy labels granule boundaries without perturbing endogenous PHB synthesis or growth^37^ (Fig. 1B, D). Phase and fluorescence images were acquired with minimal delay, and lineages were segmented using the DELTA deep-learning pipeline^38^ (Fig. 1C). This approach allows us to follow growth, cell size, and PHB dynamics for tens of thousands of lineages over week-long experiments, including multiple programmed medium switches.

### Baseline growth and PHB distributions

Under carbon- and nitrogen-replete conditions, *C. necator* grew robustly in trenches for >10 generations with mean doubling times of 2.5 ± 0.18 h. Size distributions were likewise comparable, with a mean area of 2.53 ± 0.04μm². PHB granules were visible in nearly all wild-type cell lineages, typically numbering 1–2 per cell with diameters in the hundreds of nanometers. Granule size and abundance distributions were stable over time, providing a baseline against which responses to nutrient perturbations can be assessed. In the stationary phase, *C. necator* cells are typically smaller than during exponential growth, and their morphology varies with the starvation regime. Under very low nutrient concentrations, the *C. necator* cells close to the main channel would exhibit morphology similar to that of the exponential growth phase, while cells underneath exhibit starvation morphology.

## Single-cell inheritance generates storage-rich and storage-poor lineages without growth penalty in replete conditions

To establish a baseline for PHB organization in *C. necator*, we first quantified granule presence and inheritance in steady replete conditions. At the population snapshot, cells were nearly binary: most carried either no detectable granule or a single dominant granule, with rare cases of two smaller granules (Fig. 2a).

**Figure 2.**
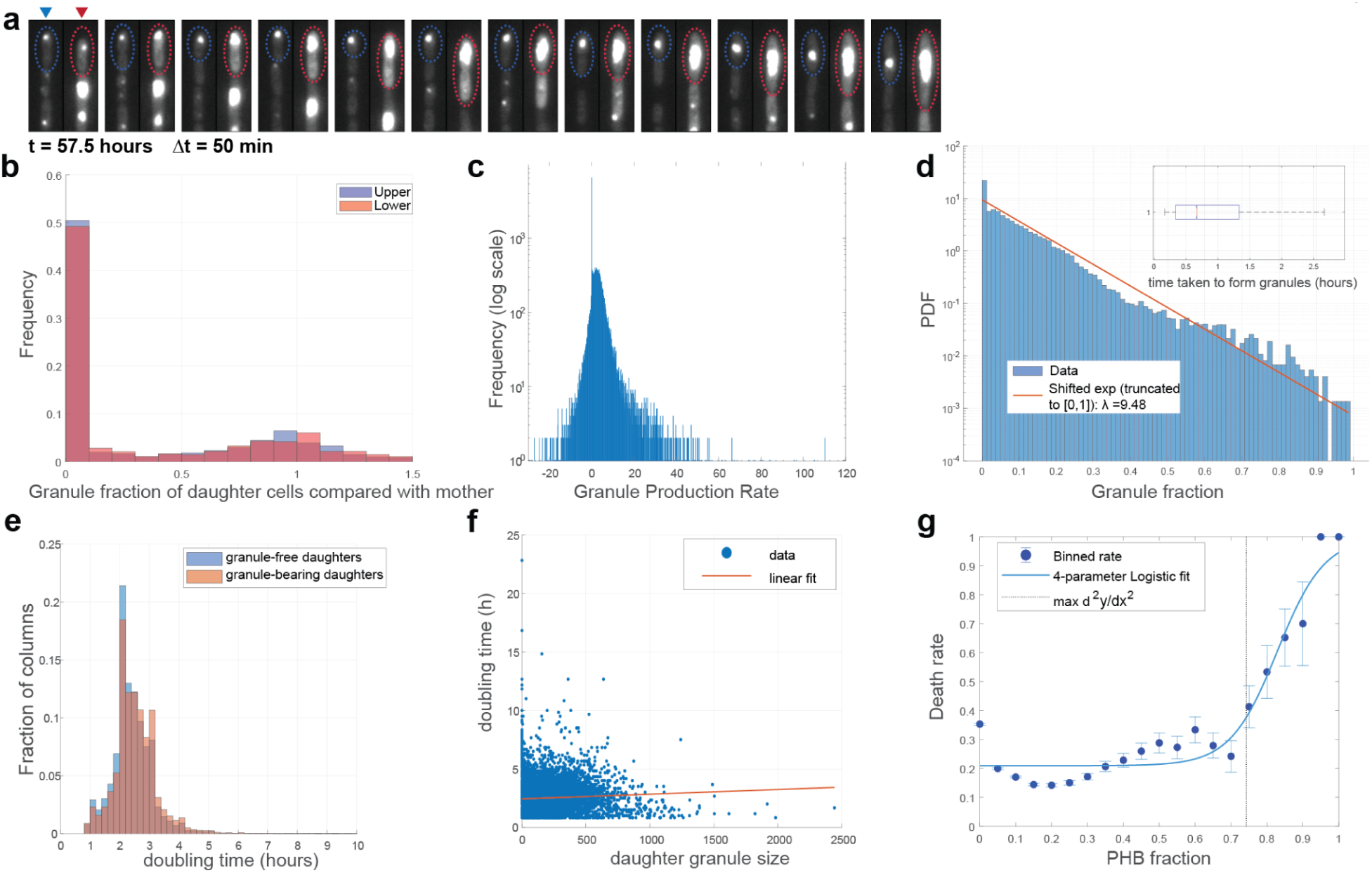
All-or-nothing inheritance of PHB granules generates storage-rich and storage-poor lineages without growth cost in replete conditions. **a.** Division-level traces showing complete inheritance of the parental granule by one daughter while the sibling receives none. This inheritance is independent of trench position. **b.** Histogram of granule fraction per daughter cells compared with their mother(0 - 1). Data from n = 46095 cells in one experiment. **c.** Granule production-rate distribution with zero-inflated/hurdle fit. Zeros arise from cells without granules. Data from n = 32313 cells in one experiment. **d.** Distribution of granule sizes among granule-bearing cells. Survival of time-to-first-granule is inset. Data from n = 59972 cells in one experiment. **e.** Doubling-time distributions of granule-bearing vs granule-free daughters. Data from n = 59972 cells in one experiment. **f.** Growth rate as a function of granule size, showing no correlation across the typical size range. Data from n = 59972 cells in one experiment. **g.** Logistic regression of arrest probability vs storage fraction, revealing a threshold penalty only at extreme granule sizes. Data from n = 59972 cells in one experiment.

Surprisingly, single-cell tracking revealed a strict, all-or-nothing inheritance rule at cell division. When a parent carried a granule, one daughter consistently inherited the entire granule while the sibling received none, with ∼50:50 probability (Fig. 2b). This partitioning was independent of trench position and channel flow. This partition rule naturally explains the zero-inflation seen in per-cell production-rate histograms: cells without a granule cannot increase granule size and therefore contribute zeros (Fig. 2c). Among granule-bearing cells, production rates followed a continuous unimodal distribution.

This all-or-nothing segregation dictates the broad yet stable granule size distribution across lineages (Fig. 2d). Cells starting without a granule typically acquired one within less than a cell cycle (median Tacq ≈ 40 mins, Fig. 2d inset), ensuring a steady fraction of granule-bearing cells. Once acquired, a granule grows over successive divisions until it is stochastically lost to a sibling. Under this rule, the probability of retaining a granule for k generations falls off geometrically (∼2−k), meaning older, larger granules become progressively rarer. A minimal segregation–growth model confirms that the combination of continuous intra-lineage growth and geometric inter-lineage loss yields the exact stationary, shifted-exponential distribution we observe (Supplementary Fig. S5).

We next examined whether carrying a granule influenced growth. Doubling times of granule-bearing vs granule-free daughters immediately after division were indistinguishable (Δμ ≈ -0.12 h, 95% CI [-0.11, -0.13]; Fig. 2e). Although the difference between doubling times is statistically distinguishable given our large sample size, this effect is negligible relative to cell-cycle variability. Granule size was likewise uncorrelated with growth rate across the typical range (slope ≈ 0.0004, Fig. 2f).

A rare penalty emerged only at extreme storage levels. Cells carrying very large granules had a sharply increased probability of division arrest (Fig. 2g). Logistic regression of arrest vs storage fraction identified an inflection point near 74% of cell area/volume.

Together, these results show that in a nutrient-replete steady state, PHB is organized by stochastic all-or-nothing inheritance, generating storage-rich and storage-poor lineages. This inheritance rule explains the zero inflation in production data and yields a geometric-like presence distribution. Crucially, PHB carriage is growth-neutral under abundance, with a rare cost only at extreme storage loads, representing only ∼0.19% of the total population. This baseline framework sets the stage for our later analyses of PHB regulation during nutrient shifts and fitness consequences at starvation boundaries(fig. 4gh).

## Replete perturbations reveal perfect adaptation of PHB fraction with asymmetric transients

We investigated whether PHB levels are actively regulated rather than passively accumulated. Baseline distributions at two replete nutrient levels (higher and lower C/N) were indistinguishable, hinting at regulation toward a common state. To test whether *C. necator* populations return to this setpoint after sustained nutrient shifts within growth-permissive regimes, we performed perturbation experiments in the microfluidic platform.

To address this, we performed perturbation experiments in the microfluidic platform. Stationary-phase cultures (40–48 h) were loaded into mother-machine chips, where replicate lanes were supplied with independent medium flows. Medium exchanges were executed with a pinch-valve manifold at <2 min per switch, faster than a cell cycle, allowing sharp perturbations without confounding by slow exchange. In two independent experiments with different initial conditions and nitrogen concentrations, the population-average PHB fraction diverged during wake-up but converged to the same steady value by ∼30 h (Fig. 3a, dashed line). Comparing size distributions, PHB fractions that were sharply different at 10 h (two-sample test, P = 2.3 × 10⁻²⁴) became indistinguishable by 30 h (Fig. 3b, top). Cell length at division and doubling-time distributions likewise converged across all conditions (Fig. 3b, middle and bottom). Returning to the same steady PHB fraction and growth distributions after a sustained change fulfills the operational integral control theory definition of perfect adaptation and is consistent with regulation to a setpoint in replete conditions.

**Figure 3.**
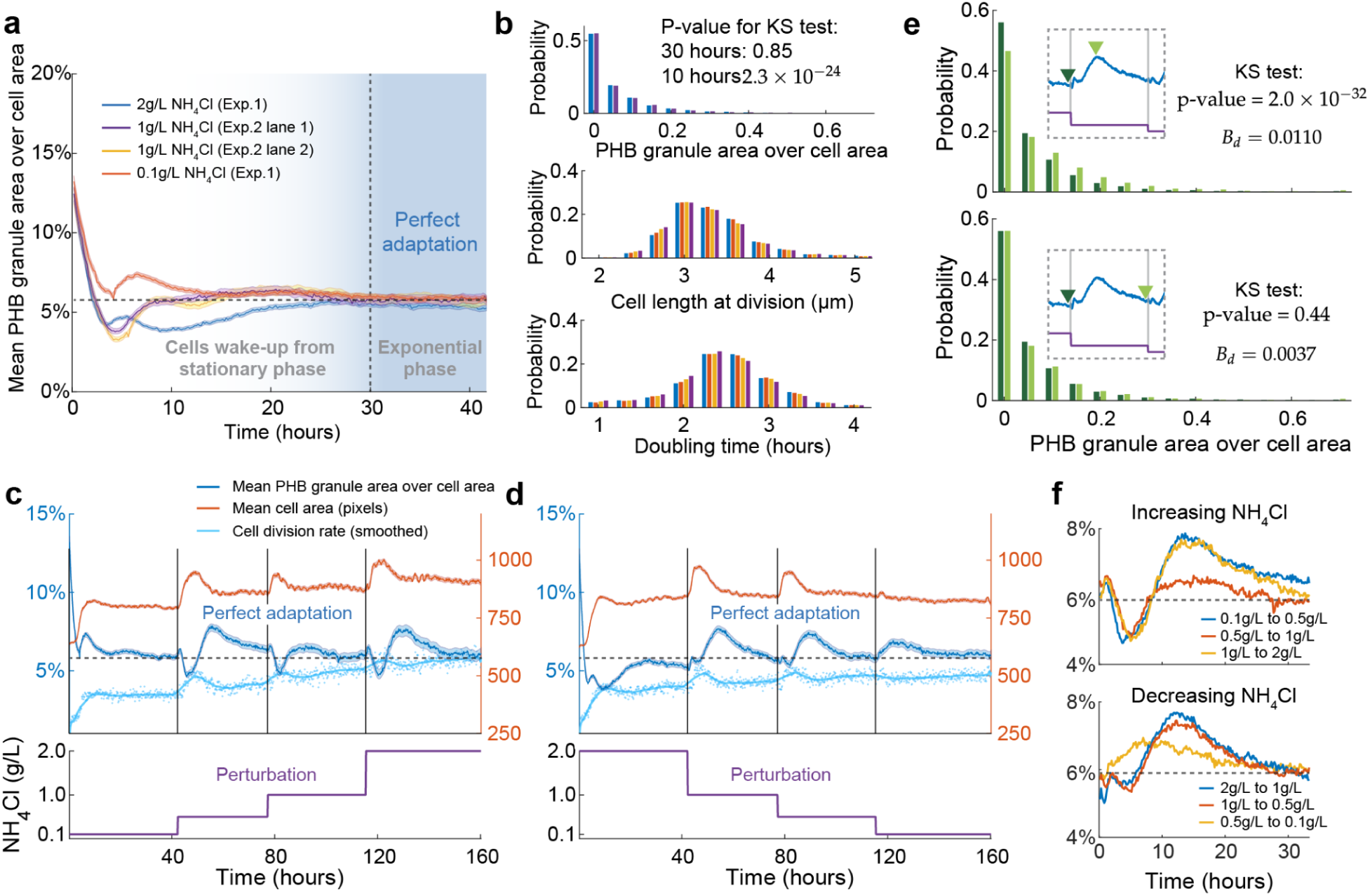
PHB fractions exhibit perfect adaptation across nutrient perturbations. **a.** Recovery from distinct initial states: stationary-phase cultures with different pre-growth histories fed replete media differing in NH₄Cl concentration. Despite divergent “wake-up” trajectories, population-average PHB fractions converge to the same steady value (dashed baseline). Traces are from two independent experiment runs (marked as Exp. 1 and Exp. 2 in the panel), with n = 3500+ cells per timepoint per trace. **b.** Distribution-level adaptation: PHB fractions significantly different at 10 h (P = 2.3 × 10⁻²⁴) converge by 30 h (top). Cell length at division (middle) and doubling time (bottom) distributions likewise converge across conditions. Data are from n = 22364 cell cycles in one experiment. **c-d.** Step perturbations in NH₄Cl spanning 200-fold (<0.01–2 g L⁻¹) produce transient deviations followed by return to baseline with near-zero steady-state error. Traces in either panel are from one independent experiment run, with n = 3500+ cells per timepoint per trace. **e.** Snapshot distributions show distinct peak response states but overlapping pre- and post-adaptation distributions. Traces in either panel are from one experiment run, with n = 3500+ cells per timepoint per trace. **f.** Asymmetric responses: step-ups produce dip/overshoot; step-downs produce elevation/decay. Second step-up (0.5 → 1 g L⁻¹) shows muted overshoot, consistent with refractory dynamics. Attenuated responses appear at the lowest NH₄Cl steps. Traces are from one experiment run, with n = 3500+ cells per timepoint per trace.

We then imposed step perturbations in nitrogen while remaining in non-limiting regimes. Across a ∼200-fold NH₄Cl range (<0.01 to 2 g L⁻¹), PHB fraction deviated transiently but returned to the same baseline with near-zero steady-state error (with a 95% CI of [0.0555, 0.0562]) (Fig. 3c–d). Distributional snapshots confirmed that pre- and post-perturbation states were indistinguishable, while peak responses differed (Fig. 3e). Transients were asymmetric: step-ups produced a dip, overshoot, and return; step-downs produced an elevation and decay (Fig. 3f). A second up-step (0.5 → 1 g L⁻¹) showed a muted overshoot, suggesting refractory behavior on the order of tens of hours. At very low nitrogen (0.1 → 0.01 g L⁻¹), response amplitude diminished, consistent with limited actuation at the edge of growth permissibility. Analogous behavior was observed for carbon perturbations. By contrast, control switches in which nitrogen was unchanged, or NaCl was added to alter ionic strength, produced no response, confirming that the dynamics are nutrient-specific rather than artifacts of switching.

A common intuition is that PHB granules buffer growth against input fluctuations. Several observations argue against this explanation. Perfect adaptation requires return to the same PHB fraction after sustained steps, which buffering alone would not enforce. Growth distributions are indistinguishable between cells with and without granules, and ΔphaC mutants grow like WT in replete conditions (Fig. 2e inset). When measured, ΔphaC populations also showed convergence of growth-rate distributions after nutrient steps, demonstrating that constancy of growth dynamics does not require PHB stores. These results favor an active regulatory loop maintaining a PHB and growth setpoint in replete regimes over a passive buffering model using PHB as a carbon ‘capacitor’. Alternatively, this setpoint could be constrained by biophysical feedback rather than purely metabolic sensing; for instance, the sheer physical volume of the granules may induce macromolecular crowding effects, altering intracellular concentrations and diffusion dynamics in ways that mechanically or spatially arrest further polymer synthesis.

Phenomenologically, the signatures of this system—zero steady-state error across ∼200-fold input variation, distributional convergence from distinct starting states, asymmetric transients, muted overshoot on repeated perturbations, and loss of control at very low nutrient—are all consistent phenomenologically with integral feedback control, a well-established mechanism for robust perfect adaptation in biological systems^39,40^. However, we do not distinguish integral feedback from other mechanisms that can produce perfect adaptation at the phenomenological level. Importantly, ΔphaC strains also show adaptation, arguing against PHB buffering as the primary mechanism. In ecological terms, regulating growth and storage toward a constant setpoint, rather than scaling directly with instantaneous nutrient supply, may be advantageous in the fluctuating environment of soil where over-responding to transient pulses is costly. Because the relationship between nutrient availability and growth rate is fundamentally concave, Jensen’s inequality dictates that a fluctuating growth rate yields a lower time-averaged population expansion than a steady growth rate. Thus, maintaining perfect adaptation via integral feedback effectively filters out high-frequency noise, maximizing long-term exponential yield. In the following section, we show that once inputs cross starvation thresholds this regulation degrades, and PHB itself confers decisive fitness benefits.

## PHB confers fitness advantages at starvation boundaries

In the previous section, we showed that *C. necator* regulates PHB fractions toward a common setpoint across a wide nutrient range, consistent with integral feedback control. We next asked how this regulation breaks down once nutrients fall below the replete regime, and whether PHB itself provides direct fitness advantages at starvation boundaries.

When nitrogen levels were reduced to ultra-low concentrations, single-cell trenches displayed a striking separation between non-growing, PHB-accumulating cells upstream and dividing cells downstream, reflecting the gradient of NH₄Cl across the trench from ∼10⁻⁶ g/L at the inflow to below 10⁻⁸ g/L at the distal end (Fig. 4a). Contrary to the expectation that extremely high C/N ratios might trigger a high-PHB mode, we found the opposite: PHB production fell sharply once nitrogen dropped below 0.01 g/L, while division rates remained unchanged (Fig. 4b–c). For example, at 0.01 and 0.001 g/L NH₄Cl, doubling times were 2.50 ± 1.08 and 2.59 ± 2.06 h, respectively, with no significant difference (P = 0.2). These results indicate that cells under severe nitrogen limitation prioritize allocation of nitrogen to growth over PHB synthesis, suppressing storage rather than enhancing it. This finding contrasts with industrial paradigms where high C/N drives bulk PHB accumulation, and aligns with recent arguments that nitrogen scarcity drives prioritization of translational capacity over storage^30,32^.

**Figure 4.**
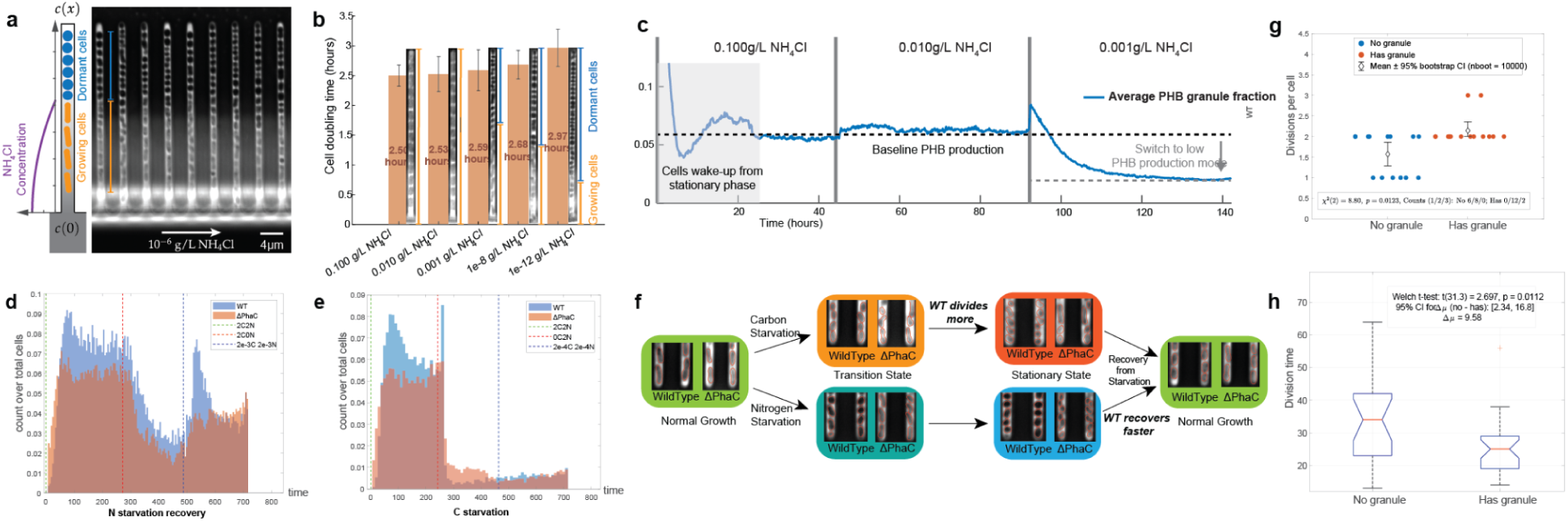
Starvation thresholds and single-cell assays. **a.** Trench gradients under ultra-low nitrogen reveal non-growing, PHB-accumulating cells upstream and dividing cells downstream (2 g/L acetate; inflow NH₄Cl = 10⁻⁶ g/L, trench minima <10⁻⁸ g/L). **b.** Doubling times (means ± SD, n=20/condition) are unchanged across 0.1, 0.01, 0.001, 10⁻⁸, and 10⁻¹² g/L NH₄Cl; e.g., 0.01 vs 0.001 g/L = 2.50 ± 1.08 vs 2.59 ± 2.06 h, P = 0.2. Data are from 20 cells (n = 20) in one independent microfluidic experiment per concentration. Data from n = 10 cells per concentration. **c.** PHB fraction drops sharply below 0.01 g/L NH₄Cl, indicating suppression of storage under extreme nitrogen limitation. Data are from one independent microfluidic experiment. Data from n = 10000+ cells per time point. **d.** Mother-machine trajectories show WT resuming divisions more rapidly than ΔphaC after nitrogen return. Data are from one independent microfluidic experiment. Data from n = 67827 cells in one experiment. **e.** Under carbon withdrawal, WT lineages yield ∼30% more progeny before arrest compared to ΔphaC. Data are from one independent microfluidic experiment. Data from n = 40224 cells in one experiment. **f.** Phenotypic summary: under carbon starvation, both strains become small and round with no PHB; under nitrogen starvation, WT retains PHB-filled cells while ΔphaC does not. **g.** Comparison of division numbers between cells with and without a granule under carbon starvation. Applying a chi-square test of independence to the two groups yielded χ²(2) = 8.80, p = 0.012, indicating that division frequency depends on granule presence. Data from 30 cells. **h.** Comparison of the time between division events in cells with and without a granule under carbon starvation. Applying a Welch t-test to the two groups yielded t(31.3) = 2.697, p = 0.0112, indicating that there’s a significant difference between the division periods in the two groups. Data from 30 cells.

Under carbon withdrawal, both wild-type and ΔphaC strains eventually ceased dividing, but wild-type lineages consistently produced ∼30% more progeny before arrest (Fig. 4e). Both strains became small and round with no detectable PHB at the end of the carbon-off interval (Fig. 4f), consistent with complete mobilization of the polymer. In our comparison of division numbers between cells with and without a granule under carbon starvation, we applied a chi-square test of independence to the two groups(Fig. 4g), and the result reveals granule-bearing cells divided more frequently and with shorter division intervals under carbon starvation (χ² test, p=0.012; Welch’s t-test, p=0.011)(Fig. 4h), indicating that there’s a significant difference between the division periods in the two groups. These observations indicate that PHB accumulated during prior replete growth is mobilized to support additional cell divisions after external carbon is exhausted, effectively increasing reproductive output.

In nitrogen starvation, the advantage was expressed at recovery rather than entry. To quantify the length of the lag phase in each group, we have implemented a piecewise regression model with an initial flat segment fitting the lag phase, followed by an exponential segment fitting the log growth phase. When NH₄Cl was restored after a period of deprivation, wild-type populations resumed exponential growth with almost no lag. In contrast, ΔphaC strains consistently exhibited a ∼2 h delay before reinitiating division (Fig. 5a). The lag difference was evident both in flow-cytometry time courses and in mother-machine trajectories, where wild-type growth was well described by the exponential fit. At a growth rate of ∼0.3 h⁻¹, this 2 h head start translates into roughly a 20–25% fitness advantage per recovery pulse, equivalent to one additional round of doubling during repeated cycles of feast and famine. These data suggest that PHB granules accumulated during nitrogen limitation serve as an immediately accessible internal reserve that enables wild-type cells to re-enter exponential growth more rapidly once nutrients return.

**Figure 5.**
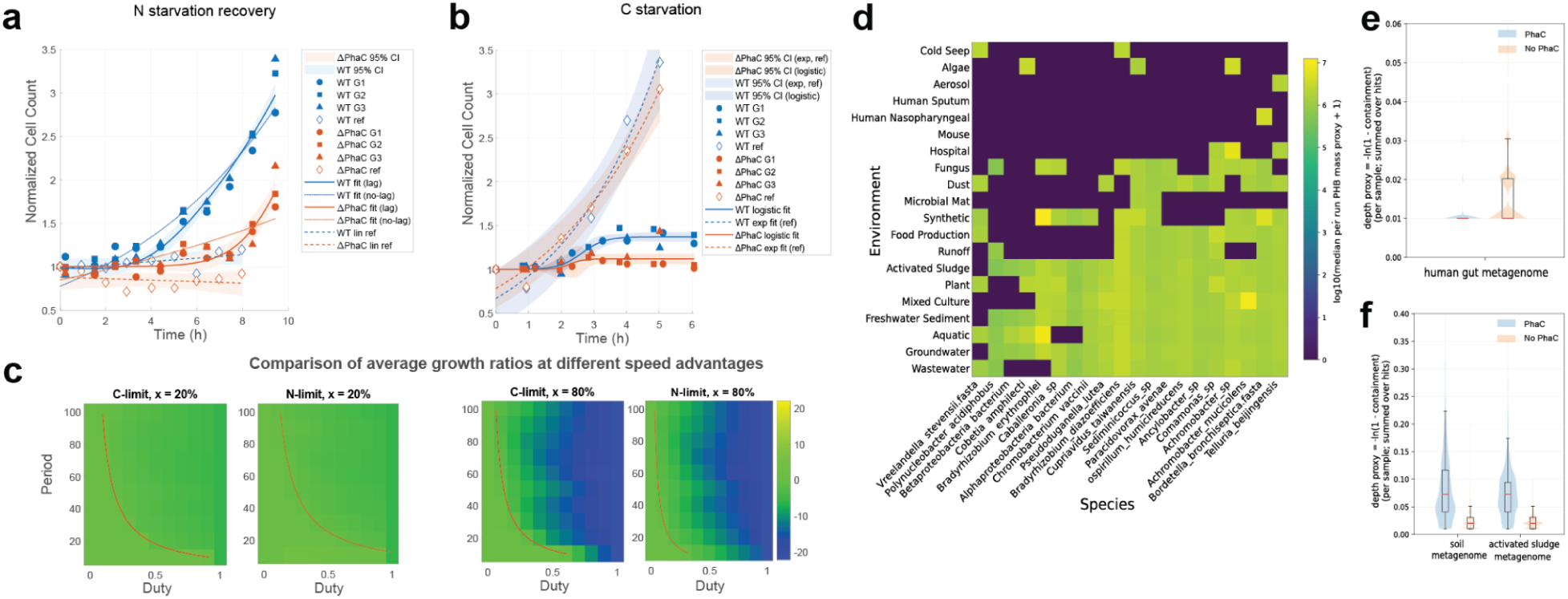
Population recovery and ecological modeling. **a.** Flow-cytometry recovery curves after nitrogen return: WT resumes exponential growth with minimal lag, ΔphaC requires ∼2 h lag. Fits: exponential (WT) vs exponential-with-lag (ΔphaC). Data represent n = 3 independent biological replicates. **b.** Offspring yield during carbon withdrawal: WT ∼30% higher than ΔphaC (mean ± CI). **c.** Pulsed-environment model integrating entry and exit advantages. The color in each heatmap indicates the competitive advantage of *C. necator* (CN) relative to the competitor (PA), quantified as the log ratio of final abundances, ln(N_CN,final / N_PA,final). Lighter colors correspond to a larger advantage of CN. The red curve marks parameter combinations where the final abundances are equal (ln ratio = 0), computed from the analytic equal-fitness condition (Supplementary Note S6). WT outcompetes a non-storer whenever duty cycle < 0.3 or period < 12 h, even when the competitor grows up to 80% faster in steady state. **d.** Heatmap of PHB-associated microbial abundance across environments. Rows denote environments and columns denote PHB-associated microbial species; values are the median estimated abundance per environment. For visualization, we selected 20 environments from the top 100 ranked environments and 20 species from the top 50 ranked strains (Methods). Abundance was estimated from genome coverage in environmental metagenomes queried with Branchwater. Two-dimensional hierarchical clustering of the environment-by-species matrix reveals two broad environment groups that differ in the diversity and composition of PHB-associated organisms. **e-f.** Box (median/IQR) and violin (density) plots of 38 PHB-positive and 38 PHB-negative species in the class Betaproteobacter per human gut metagenome bioproject (e), soil metagenome bioproject (f), and activated sludge metagenome bioproject (g). PHB-negative species are much more abundant in human gut samples, while scarce in soil metagenome samples. PHB-positive species are relatively depleted in human gut samples, while abundant in soil metagenome samples.

To assess the potential benefits of PHB storage in oligotrophic, competitive settings, we built a simple two-strain model comparing C. necator (CN) to a non-storing competitor (PA) under a pulsed nutrient regime spanning duty cycles and periods (Fig. 5c). Each cycle consisted of an abundance phase of duration D·P and a starvation phase of duration (1−D)·P, during which either carbon or nitrogen was removed. Duty cycle (D) was defined as the fraction of time with abundant nutrients, and period (P) as the total cycle length. During abundance, both strains grew exponentially with strain-specific rates (r_CN for CN and r_PA = r_CN·(1 + x/100) for PA, where x is the competitor’s percent speed advantage).

Under carbon starvation, PA ceased growth immediately, whereas CN received a one-time 1.3-fold increase in abundance at the onset of starvation (implemented as an instantaneous fold-change to represent additional progeny production prior to arrest). Upon return to complete medium, both strains underwent a transient recovery phase implemented as reduced-rate growth (half the basal rate) for a fixed duration before resuming full exponential growth. Under nitrogen starvation, both strains stopped dividing; after nitrogen was restored, PA experienced a one-generation no-division lag before resuming exponential growth, whereas CN resumed growth immediately.

We summarized outcomes as heatmaps in which color encodes the growth advantage of CN relative to PA, quantified as the natural-log ratio of final abundances ln(N_CN/N_PA). Lighter colors indicate a larger advantage, and the overlaid red curve marks parameter combinations where both populations reach the same final abundance (zero log ratio), computed from an analytic equal-fitness condition (Supplementary Note S6). Model details and heatmap construction are described in the Methods.

Here, C. necator markedly outperformed the competitor under short duty cycles and brief periods, allowing it to compete effectively even against a competitor with steady state growth rates up to 80% higher. Although highly simplified, these parameter ranges are broadly consistent with pulse-driven soil environments, where microbial access to carbon and nutrients is shaped by episodic wetting and localized inputs from roots and rhizodeposition^41,42^. Storage thus acts as a conditional fitness strategy: costly under long, stable abundance, but strongly favored under short, fluctuating regimes.

Motivated by our microfluidic results, we asked whether this physiological mechanism manifests as a recognizable ecological pattern. By matching curated genomes of PHB-associated microbes to environmental metagenomes, we found that samples clustered hierarchically based on environmental stability (Fig. 5d, Fig. S4). Variable habitats (both natural and engineered) harbored diverse and abundant PHB-associated species, whereas stable host-associated environments (e.g., human guts) contained far fewer.

To perform a more controlled comparison, we contrasted the distribution of closely related *Betaproteobacteria* species with (n=38) and without (n=38) the *phaC* gene across distinct stability regimes. Consistent with our hypothesis, we observed strong environmental stratification (Fig. 5e-g). The non-PHB group dominated the host-buffered gut microbiome. In contrast, the PHB-storing group was enriched in stochastic soils and reached maximal abundance in the engineered feast-famine cycles of activated sludge.

Taken together, these results challenge the conventional view of PHB as a generic overflow response to high C/N ratios^6,14^. Instead, storage emerges as a state-dependent allocation strategy: suppressed when nitrogen is limiting, neutral in steady replete conditions, and decisive at starvation boundaries, where it extends progeny yield during carbon withdrawal and accelerates regrowth during nitrogen replenishment. By integrating these advantages over repeated pulses, PHB enables *C. necator* to persist and compete effectively in oligotrophic, fluctuation-dominated ecosystems, a niche where canonical “fast growers” are disadvantaged by their dependence on stable abundance.

Our results reveal a simple organizing principle for PHB in C. necator: neutral in replete steady state, decisive at starvation boundaries. As nutrient inputs drop below the replete regime and setpoint control degrades, the mobilization of storage becomes directly beneficial. This dynamic parallels Alon’s “last generation” observations in *E. coli*, in which entry into the stationary phase is preceded by a transient burst of nutrient-uptake and stress response followed by an abrupt growth arrest^43^. This state-dependent utility is consistent with the view that intracellular carbon storage provides a fitness buffer in feast–famine environments, allowing cells to invest during resource pulses and sustain growth and survival during subsequent scarcity.^2,6,44^ Indeed, the ecological importance of this dynamic is further supported by a recent bulk-culture study in the marine Alphaproteobacterium *Phaeobacter inhibens*, demonstrating that PHB is consumed during carbon starvation and that ΔphaC mutants exhibit significantly reduced long-term viability^45^. Alongside analogous observations for glycogen storage^44^, this independent evidence from a completely different phylogenetic lineage and ecosystem (marine vs. soil) underscores that intracellular carbon storage is a broadly conserved survival strategy.

Crucially, the reproductive dividend realized during carbon starvation provides an evolutionary rationale for the all-or-nothing inheritance rule established in replete conditions. Why would a population adopt such stark asymmetric partitioning? We propose that this strategy solves the problem of resource indivisibility at the single-cell limit. Because cell division imposes a discrete energetic threshold, a symmetric distribution of finite PHB stores during sudden starvation could dilute the reserves of both daughter cells below the minimum required to complete a cycle, yielding zero additional progeny. By concentrating storage into a single lineage, *C. necator* ensures that at least one subpopulation cleanly crosses the division threshold. In this light, the all-or-nothing partitioning is not a regulatory failure, but a strategy that prevents lethal resource dilution and mathematically guarantees a population-level dividend under severe starvation.

While the industrial paradigm for *C. necator* posits that a high carbon-to-nitrogen (C:N) ratio acts as a metabolic toggle for poly(3-hydroxybutyrate) (PHB) accumulation^23,24^, our single-cell observations suggest this relationship is not a simple linear function of nutrient stoichiometry. In bulk fed-batch systems, the ’high-PHB mode’ is typically a transient response to acute nitrogen exhaustion. In contrast, our data from consistent oligotrophic environments reveal an alternative allocation logic: when nitrogen is limiting but persistently available, the organism appears to prioritize the maintenance of growth machinery over carbon sequestration. This suggests that *C. necator* employs a behavior consistent with an integral feedback mechanism to stabilize the specific growth rate, only pivoting to massive PHB accumulation when a total depletion of nitrogen triggers the response. This helps reconcile why bulk strategies—which are optimized for total yield following nutrient exhaustion —differ from single-cell behaviors in stable, nutrient-lean regimes, highlighting how the same metabolic network can implement distinct survival strategies across different regions of the nutrient landscape.

This dynamic allocation strategy maps onto the conditions of pulse-driven soils, where rainfall and root exudation generate short, intermittent resource windows. In such erratic environments, the perfect adaptation we observed during replete growth acts as a critical filter. By resetting physiology toward a common baseline instead of tracking each transient fluctuation, it buffers cells against overcommitting to short, noisy pulses and reduces the metabolic penalty of frequent adjustments. In this view, storage is a state-dependent strategy that is neutral during stability but wins the transitions that structure life in oligotrophic landscapes.

Qualitative patterns from our metagenomic survey align with this prediction: the marked enrichment of storage-capable species in fluctuating habitats supports the view that the “loop-breaking” fluctuations we simulated in the laboratory are not merely artificial stressors, but represent a fundamental selective filter in nature. The prevalence of this high-investment strategy in variable environments underscores that for bacteria like *C. necator*, the ability to navigate frequent entry and exit from starvation is a defining ecological competence.

Several limitations and alternative explanations merit mention. Firstly, we treat integral feedback as a phenomenological description of replete dynamics; we do not identify molecular components. Known PHB regulators (e.g. phasins^46^, PhaR-like DNA binding proteins^47^, PhaZ PHB depolymerases^48^) and global C/N sensors could implement parts of this loop, but distinguishing transcriptional, metabolic, and post-translational contributions will require targeted perturbations (e.g., *phaR* mutants, inducible depolymerase, nitrogen-sensing knockdowns) and time-resolved expression/flux measurements. Secondly, our granule labeling uses a catalytically inactive PhaC1–EYFP under a low-copy plasmid and native promoter; while growth and PHB dynamics match unlabeled controls within variance, subtle tag-specific effects cannot be fully excluded. Third, gradients within trenches at ultra-low nitrogen are intrinsic to diffusion-limited supply; we leveraged them to bracket thresholds, but future designs with shorter trenches or controlled perfusion could sharpen those measurements. Finally, while ΔphaC provides a clean genetic contrast, other storage pathways (e.g., glycogen) may contribute under some conditions; the present work isolates PHB’s role in *C. necator* but does not exclude auxiliary reserves in other taxa. Furthermore, our metagenomic analysis is qualitative and correlative.

The complexity of natural communities and confounding factors—such as the drastically larger genomes of *phaC*-possessing lineages—prevent attribution of ecological success solely to the PHB pathway. This analysis also relies on the assumptions that *phaC* presence is a robust proxy for storage capacity and that broad environmental categories consistently reflect local nutrient fluctuation regimes. All experiments were performed at 30°C in defined media; PHB regulation may differ under other carbon sources, temperatures, or community contexts.

This framework leads to testable predictions. Replete-regime adaptation metrics (steady-state error, adaptation time, overshoot) should be invariant to carbon/nitrogen steps within the permissive window but degrade abruptly as cells approach starvation thresholds. Entry advantage should scale with pre-starvation PHB load and vanish in ΔphaC or depolymerase-overexpressing strains that prematurely deplete PHB; exit advantage should scale with the nitrogen-free interval and diminish when PHB cannot be mobilized. Ecologically, storage-rich genotypes should be enriched in pulse-dominated environments (short duty, short period) even when slower in steady abundance, whereas non-storers should dominate under long, stable nutrient availability.

Beyond ecology, these insights suggest biotechnological levers. If setpoint regulation acts upstream of granule synthesis, manipulating the controller (not just PhaC activity) could stabilize yields across feed fluctuations in industrial PHB processes. Conversely, to maximize recovery after process upsets, tuning depolymerase or mobilization pathways may shorten lags without compromising steady-state performance.

In sum, *C. necator* integrates robust regulation in replete regimes with conditional storage payoffs at starvation boundaries. This resolves apparent contradictions between industrial PHB paradigms and single-cell behavior in oligotrophic conditions and offers a quantitative basis for predicting when—and why—storage strategies are favored.

## Method

### Bacterial Strains

The EYFP marked strains are *R. eutropha H16* with plasmid

*pBBR1MCS-2-PhaC-phaC1-C319A-eyfp* from Pfeiffer Lab^37^, a universal vector for construction of fusions C terminal to EYFP under the control of the phaCAB promoter. The PhaC knockout strains are donated from the Criddle lab at Stanford University.

### Growth condition, microscopy, and microfluidics

Cultures were grown overnight in M9 minimal medium with 2g/L NaAc and 0.5g/L NH4Cl at 30°C. At OD600 = 1.0, the cell culture was 10x concentrated by centrifugation for injection into the mother machine. The cells were loaded by centrifugation at 850g for 90 seconds.

Fresh M9 medium was infused by a syringe pump. For each experiment, images were acquired from 60 – 80 fields at ten-minute intervals using ZEN software and a Zeiss AXIO Observer 7 Inverted LED Fluorescence Microscope equipped with a motorized stage and an Axiocam 712 mono camera. The standard LED TL lamp was selected for the white light source used for phase imaging, and the Colibri 7 Multicolor LED Light Source was selected for fluorescence imaging. Our device consists of 12000 growth channels and an automated microscope stage, which we use to continuously scan 60-80 fields of view for over 150 hours at ten-minute intervals. Since each field of view contains ∼1000 cells at 63x magnification, we acquire and process images of ∼1e07– 1e08 individual cells per experiment. See Supplementary Experimental Procedures for more details.

### Image analysis and data acquisition

DeLTA^38^ was used to analyze time-lapse images. We analyzed typically ∼1e07 cells per each time-lapse experiment. The current version of software can complete analysis in about 8-10 hours on an RTX 3080Ti and Intel Core i9-12900KF workstation with 128GB memory.

Segmentation of cells was performed by the U-Net segmentation model in DeLTA.

### Flow cytometry and corresponding data analysis

WT and PhaC knockout (ΔPhaC) *C. necator* cells were cultured in M9 medium to stationary phase, then back-diluted and cultured to exponential growth phase. Cells were then normalized to a similar OD, and proceeded with further experiments. In carbon starvation experiments, cells were spun down at 1300g for 90 seconds, then transferred to M9 minimal medium either with 0g/L NaAc and 2g/L NH_4_Cl or with 2g/L NaAc and 0.5g/L NH_4_Cl, for carbon starvation and no starvation control respectively. For nitrogen starvation recovery experiments, cells were first cultured overnight in M9 minimal medium with 2g/L NaAc and 0g/L NH_4_Cl, then transferred to M9 minimal medium either with 2g/L NaAc and 0.5g/L NH_4_Cl for recovery, or with 2g/L NaAc and 0g/L NH_4_Cl for no recovery control. The cells were then washed twice with the respective medium. Flow cytometry measurement of cell density was performed every hour by a Sony SH800S cell sorter. Samples from the respective cultures were diluted 1:10, then fed into the flow cytometer for 90 seconds. Cell count was collected for the middle 30 seconds, after the count per second reading settled down.

After obtaining the cell count data, all cell counts were normalized with respect to the first time point. Then logistic regression was used on carbon starved cells, while the no starve control was fitted to an exponential growth model. On the nitrogen recovery side, a linear model was used to fit the no recovery control, while the normalized cell count of recovering cells were fitted to a piecewise model with a flat segment fitting the lag phase, and an exponential model fitting the exponential growth phase.

### Pulsed-regime simulations and analytic equal-fitness contours

We simulated repeated feast–famine cycles with period P and duty cycle D using a binary nutrient-availability signal R(t) that equals 1 during the abundance interval and 0 during the starvation interval. Specifically, R(t)=1 when mod(t,P) < D*P and R(t)=0 otherwise, where mod(t,P) denotes time modulo the cycle period. Strain abundances b_CN(t) and b_PA(t) were updated by forward iteration with fixed time step dt. During abundance (R=1), each strain grew exponentially at a strain-specific rate r_i (CN: r_CN; PA: r_PA = r_CN*(1 + x/100)); during starvation (R=0), growth was arrested (no division), except as noted for a recovery phase in the carbon-starvation implementation below.

Carbon-starvation regime: At each abundance-to-starvation transition (R switches from 1 to 0), CN received a one-time multiplicative fold-change of 1.3, representing PHB-supported progeny production prior to arrest. At each starvation-to-abundance transition (R switches from 0 to 1), both strains entered a recovery phase implemented as reduced-rate growth at r_i/2 for the following two generations, leading to a duration t_rec,i = 2 * T_d,half = 2*(2*ln(2)/r_i) = 4*ln(2)/r_i. In the numerical implementation, this recovery timer was decremented continuously (with time step dt), so reduced-rate growth persisted for exactly t_rec,i even if the environment switched to starvation before the timer elapsed; after the timer elapsed, growth proceeded at the full rate r_i only during times with R=1.

Nitrogen-starvation regime: Both strains ceased division during starvation. Upon re-feeding (R switches from 0 to 1), PA experienced a one-generation no-division lag of duration tau_PA = ln(2)/r_PA before resuming exponential growth at r_PA, whereas CN resumed growth immediately at r_CN.

Competitive outcome and heatmaps. For each (D,P) condition, simulations were run to a fixed horizon T_end and the competitive outcome was recorded as ln[b_CN(T_end)/b_PA(T_end)] (natural logarithm). Heatmaps display this log ratio as a function of duty cycle and period.

Analytic equal-fitness contours. To summarize the equal-fitness boundary ln[b_CN(T_end)/b_PA(T_end)]=0 without relying on numerical contour interpolation on a coarse (D,P) grid, we computed a smooth analytic equal-fitness contour for each regime by equating the per-cycle multiplicative factors of CN and PA implied by the update rules. The resulting contour has the form P = (DP0)/D, where DP0 is the threshold abundance duration that solves “per-cycle CN multiplier = per-cycle PA multiplier” (Supplementary S6) In figures, we report only this smooth analytic equal-fitness contour (red line) for clarity.

(Notes on notation: ln denotes the natural logarithm; exp denotes the exponential; max(0,z) returns z if z>0 and 0 otherwise.)

### Metagenomic analysis

#### Genome Selection and Curation

To identify potential PHB-storing organisms, the amino acid sequence of the PHB synthase (*phaC*) from *Cupriavidus necator* H16 was used as a query for a BLASTp search against the NCBI non-redundant protein database. The top 5000 hits were retrieved and taxonomically collapsed into 140 representative microbial species. From this pool, we focused on the class *Betaproteobacteria* to minimize phylogenetic distance. To enable a controlled comparison, we curated two distinct functional cohorts: a study group of 38 representative species containing *phaC*, and a control group of 38 species lacking *phaC*. Selection was based on the availability of high-quality genome assemblies in RefSeq/GenBank to ensure accurate metagenomic mapping (Fig. S7).

#### Metagenomic Querying and Analysis

Metagenomic abundance profiling was performed using the Branchwater plugin (V0.4.0) for Sourmash (V0.18.0). The curated genomes were queried against the bw_k21, s=1000 metagenomics database^49,50^, using a k-mer size of 21.

Computations were executed locally on a MacBook Pro (Apple M1 Max processor).

#### Abundance Estimation

To ensure detection reliability, only alignments meeting a containment Average Nucleotide Identity (cANI) > 0.9 and a query coverage > 0.2 were retained for analysis. The sequencing depth of each genome in a given metagenomic sample, which serves as a proxy for abundance, was estimated using the genome containment breadth (C) according to the Poisson approximation:

Depth Proxy=−ln(1−C)

This metric allows for the comparison of relative abundances across diverse environmental samples.

#### Statistics and Reproducibility

Statistical analyses were performed using MATLAB R2024b, Python 3.8.8 (imaging analysis) and Python 3.9.23 (metagenomic analysis). No statistical methods were used to predetermine sample sizes; sample sizes were dictated by the tracking capacity of the microfluidic device, yielding observational n values of tens of thousands of independent single-cell lineages per condition, which provides sufficient statistical power. Data distribution was assumed to be normal, but this was not formally tested.

For comparisons of continuous variables between two groups (e.g., division intervals between granule-bearing and granule-free lineages), two-tailed Welch’s unequal variances *t*-tests were utilized. For categorical outcomes (e.g., division frequencies), Pearson’s chi-square tests of independence were applied. The exact test statistics (e.g., t-values, χ2 values), degrees of freedom, and exact P values are reported in the main text or corresponding figure legends.

Due to the extreme technical scale and week-long duration of the microfluidic experiments, independent biological replicates (defined as distinct microfluidic chip runs from independent overnight cultures) were performed where noted (e.g., Fig. 3a). For experiments involving a single microfluidic run, statistical robustness is derived from the massive parallelization of the device. The exact number of single-cell lineages analyzed (n) for each specific experiment is provided in the corresponding figure legends.

## Supplemental information

Supplemental information includes detailed experimental protocols and microfluidic chip manufacturing procedures, comprehensive descriptions of imaging acquisition and experimental methods, as well as methodologies for segmentation, tracking analysis, and modeling.

## Author Contributions

J.H., R.Y., and A.P.A. designed the research; J.H., R.Y., Y.M., and H.M performed the research; J.H., R.Y., Y.M., H.M. and A.P.A. wrote the paper. J.H. and R.Y. contributed equally to this work. A.P.A. supervised the work and aided in analysis.

## Acknowledgments

We thank Eoin Brodie for critical reading of the manuscript and valuable feedback. This material by ENIGMA- Ecosystems and Networks Integrated with Genes and Molecular Assemblies (http://enigma.lbl.gov), a Science Focus Area Program at Lawrence Berkeley National Laboratory is based upon work supported by the U.S. Department of Energy, Office of Science, Office of Biological & Environmental Research under contract number DE-AC02-05CH11231.

This work was also supported by the U.S. National Science Foundation under award number 2125069.

## Data availability

All processed data and intermediate datasets generated or analysed during this study are publicly available on Zenodo: DOI 10.5281/zenodo.19446721. Source data for all figures are provided with this paper via the same Zenodo link. Due to their exceptionally large size (∼200 TB), the raw microfluidic imaging data cannot be hosted on standard public repositories. These raw datasets are securely archived on institutional servers at UC Berkeley and will be made available by the corresponding author upon reasonable request. Transfer of the raw data may require the requester to provide physical storage devices and cover associated shipping costs.

## Code availability

All custom scripts and code used for data processing, statistical analysis, and figure generation are publicly available on Zenodo: DOI 10.5281/zenodo.19446721.

