## Supplementary Info for "Single-cell carbon storage dynamics drive conditional fitness in microbes"

### Supplementary Information

#### S1. Nutrient diffusion modeling and validation

We used a minimal diffusion–consumption model to quantify how closely the mother cell samples the bulk ammonium concentration in the microfluidic "mother-machine" device and how strongly the side trenches perturb the main-channel nutrient supply.

##### S1.1 Estimation of the volumetric uptake rate $S_v$

To estimate the uniform volumetric consumption term $S_v$, we first approximated the nitrogen incorporation rate per cell. We assumed [61]typical bacterial biomass composition[61] $CH_{1.8}O_{0.5}N_{0.2}$, corresponding to an N mass fraction

$f_N \approx 0.11$ g N per g dry weight

Published calibrations for E. coli indicate that $10^{12}$ cells correspond to 0.39 g dry[62]eight. This gives a dry mass per cell[62]

$m_{cell} = 0.39 \text{ g} / (10^{12} \text{ cells})$
$m_{cell} \approx 3.9 \times 10^{-13}$ g per cell

The nitrogen mass per cell is

$m_{N\_cell} = f_N * m_{cell}$
$m_{N\_cell} \approx 0.11 \times 3.9 \times 10^{-13} \text{ g} \approx 4.3 \times 10^{-14}$ g N per cell

Using a molar mass of nitrogen

$M_N = 14 \text{ g mol}^{-1}$

the nitrogen content per cell in moles is

$n_{N\_cell} = m_{N\_cell} / M_N$
$n_{N\_cell} \approx 4.3 \times 10^{-14} \text{ g} / 14 \text{ g mol}^{-1} \approx 3.1 \times 10^{-15}$ mol N per cell

Under our growth conditions, the doubling time was

$t_d = 2.5 \text{ h} = 2.5 \times 3600 \text{ s} = 9000 \text{ s}$

Assuming that nitrogen uptake is dominated by incorporation into new biomass during exponential growth, the mean nitrogen incorporation rate per cell is

$r_{cell} = n_{N\_cell} / t_d$
$r_{cell} \approx 3.1 \times 10^{-15} \text{ mol} / 9000 \text{ s} \approx 3.4 \times 10^{-19} \text{ mol } (\text{cell} \cdot \text{s})^{-1}$

To convert this per-cell rate into a volumetric uptake $S_v$, we assume an effective per-cell volume and a packing fraction in the trench. We take

$V_{cell} = 3.5 \times 10^{-18}$ $m^3$ per cell
$\phi = 0.56$ (dimensionless packing fraction)

The number of cells per unit volume is

cells per volume = $\phi / V_{cell}$
cells per volume $\approx 0.56 / (3.5 \times 10^{-18}$ $m^3$ per cell) $\approx 1.6 \times 10^{17}$ cells $m^{-3}$

The volumetric nitrogen uptake rate is then

$S_v = r_{cell}$ * (cells per volume)
$S_v \approx (3.4 \times 10^{-19}$ mol $(\text{cell}\cdot\text{s})^{-1}) * (1.6 \times 10^{17}$ cells $m^{-3})$
$S_v \approx 0.17$ mol $m^{-3}$ $s^{-1}$

This $S_v$ value is used in the following diffusion–consumption model.

**S1.2 One-dimensional diffusion–consumption model**

Each side trench is approximated as a one-dimensional segment of length L extending from the main flow channel at $x = L$ to a closed end at the mother cell at $x = 0$. Along this axis, ammonium concentration $c(x)$ is governed by Fick's law with the uniform volumetric consumption term $S_v$:

diffusive flux:
$J(x) = -D * dc/dx$

mass balance with uniform consumption:
$D * d^2c/dx^2 = S_v$

Here, $c(x)$ is the ammonium concentration, D is the diffusion coefficient of $NH_4^+$ in water, and $S_v$ is the effective volumetric uptake rate of the packed cells in the trench.

We impose a no-flux boundary at the closed end and a fixed concentration at the opening:

$dc/dx = 0$ at $x = 0$ (closed end, no flux)
$c = c_{bulk}$ at $x = L$ (open end, in contact with the main channel)

Solving these equations with the above boundary conditions gives a quadratic concentration profile:

$c(x) = c_{bulk} - (S_v / (2D)) * (L^2 - x^2)$

The concentration at the mother cell ($x = 0$) is therefore

$c_{end} = c(0) = c_{bulk} - S_v * L^2 / (2D)$

The concentration drop along the trench is

$\Delta_c = c_{bulk} - c_{end}$

and the relative depletion is

$$\Delta c / c_{bulk} = S_v \cdot L^2 / (2D \cdot c_{bulk})$$

We used a trench length of

$$L = 35 \times 10^{-6} \text{ m}$$

and a diffusion coefficient for $NH_4^+$ in water of

$$D = 1.98 \times 10^{-9} \text{ m}^2 \text{ s}^{-1}$$

In our experiments, the bulk ammonium chloride concentration was 0.1 g $L^{-1}$ $NH_4Cl$. With a molar mass of 53.5 g $mol^{-1}$, this corresponds to

$$c_{bulk\_mol\_per\_L} = 0.1 \text{ g L}^{-1} / 53.5 \text{ g mol}^{-1} \approx 1.87 \times 10^{-3} \text{ mol L}^{-1}$$
$$c_{bulk} = c_{bulk\_mol\_per\_L} \times 1000 \text{ L m}^{-3} \approx 1.87 \text{ mol m}^{-3}$$

Using the $S_v$ value estimated below ($S_v \approx 0.17$ mol $m^{-3}$ $s^{-1}$), we obtain

$$c_{end} \approx 1.82 \text{ mol m}^{-3}$$
$$\Delta c = c_{bulk} - c_{end} \approx 1.87 - 1.82 \approx 0.054 \text{ mol m}^{-3}$$

The relative depletion is then

$$\Delta c / c_{bulk} \approx 0.054 / 1.87 \approx 0.03$$

so the mother cell experiences approximately 97% of the bulk ammonium concentration under these conditions.

**S1.3 Device-scale ammonium budget**

We next asked whether uptake by all side trenches significantly perturbs the main-channel ammonium flux. The convective molar flux of $NH_4Cl$ in the main channel is

$$J_{main} = c_{bulk} \cdot Q$$

where Q is the volumetric flow rate. In our experiments,

$$Q = 20 \times 10^{-6} \text{ L min}^{-1}$$

Converting to liters per second,

$$Q_{L\_per\_s} = Q / 60$$
$$Q_{L\_per\_s} \approx 20 \times 10^{-6} \text{ L min}^{-1} / 60 \text{ s min}^{-1} \approx 3.3 \times 10^{-7} \text{ L s}^{-1}$$

Using $c_{bulk\_mol\_per\_L} \approx 1.87 \times 10^{-3}$ mol $L^{-1}$, the main-channel molar flux is

$$J_{main} = c_{bulk\_mol\_per\_L} \cdot Q_{L\_per\_s}$$
$$J_{main} \approx 1.87 \times 10^{-3} \text{ mol L}^{-1} \cdot 3.3 \times 10^{-7} \text{ L s}^{-1} \approx 6.2 \times 10^{-10} \text{ mol s}^{-1}$$

To estimate uptake in the side trenches, we approximate each trench cross-section as a trapezoid based on AFM measurements, with

width: $w = 1.4 \times 10^{-6}$ m
heights: $h1 = 2.0 \times 10^{-6}$ m and $h2 = 1.0 \times 10^{-6}$ m

The cross-sectional area is

$$A_{trench} = w * (h1 + h2) / 2$$
$$A_{trench} = 1.4 \times 10^{-6} \text{ m} * (2.0 \times 10^{-6} \text{ m} + 1.0 \times 10^{-6} \text{ m}) / 2$$
$$A_{trench} \approx 2.1 \times 10^{-12} \text{ m}^2$$

The volume of one trench is

$$V_{trench} = L * A_{trench}$$
$$V_{trench} \approx 35 \times 10^{-6} \text{ m} * 2.1 \times 10^{-12} \text{ m}^2 \approx 7.4 \times 10^{-17} \text{ m}^3$$

Using the volumetric uptake $S_v \approx 0.17$ mol m$^{-3}$ s$^{-1}$, the uptake rate per trench is

$$J_{trench} = S_v * V_{trench}$$
$$J_{trench} \approx 0.17 \text{ mol m}^{-3} \text{ s}^{-1} * 7.4 \times 10^{-17} \text{ m}^3 \approx 1.3 \times 10^{-17} \text{ mol s}^{-1}$$

We estimate that the device contains

$N_{trenches} \approx 2.5 \times 10^4$ side trenches

The total ammonium uptake by all trenches is therefore

$$J_{side\_total} = J_{trench} * N_{trenches}$$
$$J_{side\_total} \approx 1.3 \times 10^{-17} \text{ mol s}^{-1} * 2.5 \times 10^4 \approx 3.2 \times 10^{-13} \text{ mol s}^{-1}$$

The fraction of the main-channel ammonium flux diverted into the side trenches is

$$fraction_{side} = J_{side\_total} / J_{main}$$
$$fraction_{side} \approx 3.2 \times 10^{-13} \text{ mol s}^{-1} / 6.2 \times 10^{-10} \text{ mol s}^{-1}$$
$$fraction_{side} \approx 5.1 \times 10^{-4} \approx 0.05\%$$

In summary, under our experimental conditions, (1) the mother cell experiences ammonium levels within a few percent of the bulk concentration (approximately 97% at 0.1 g L$^{-1}$ $NH_4Cl$), and (2) ammonium uptake by cells in the side trenches extracts only a negligible fraction (~0.05%) of the ammonium supplied by the main channel. These results support treating the mother cell as effectively sampling the external nutrient environment without substantially perturbing it. All calculations were implemented in Python; the code is provided in Supplementary Code S1.

**S2. Specificity controls.**

**Supplementary Fig. 2 | PHB granule fraction is insensitive to ionic strength but shows similar transient dynamics upon carbon and nitrogen shifts.**

**a, NaCl osmolarity control.** C. necator was cultured in media containing different NaCl concentrations while keeping the carbon and nitrogen sources constant. Switching between media with different $Na^+$ and $Cl^-$ concentrations did not elicit detectable changes in the PHB granule fraction, and the time courses remained essentially flat across all conditions. This rules out simple effects of osmolarity or ionic strength as the cause of the PHB dynamics observed in the main text. Data from n = 3600+ cells per time point.

**b, Carbon shifts at constant nitrogen.** Cells were cultured at different sodium acetate (NaAc) concentrations while maintaining a constant ammonium chloride (NH4Cl) concentration. When the carbon level was changed, the PHB granule fraction exhibited transient fluctuations around the time of the medium switch but rapidly relaxed back to approximately the pre-switch level. The response closely resembles the behaviour observed when varying NH4Cl at fixed carbon in the main text (Fig. 3). Together, these data indicate that C. necator maintains an approximately constant division rate and PHB production rate over a broad range of external C/N ratios, and that the transient PHB dynamics upon medium shifts arise from changes in C/N balance rather than from differences in Na+/Cl− concentrations. Data from n = 3500+ cells per time point.

**1**

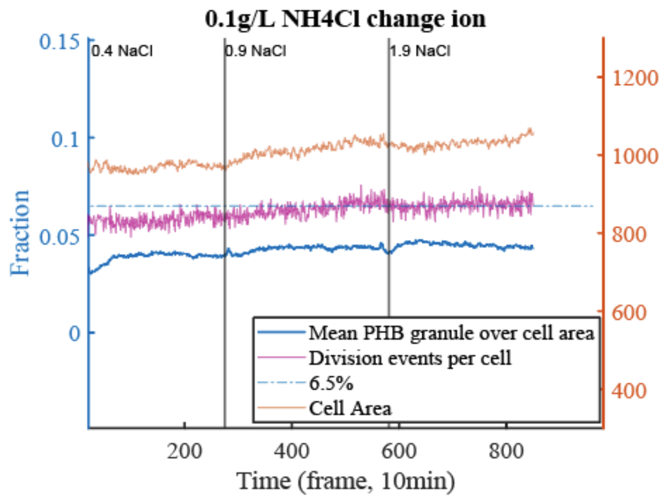


**2**

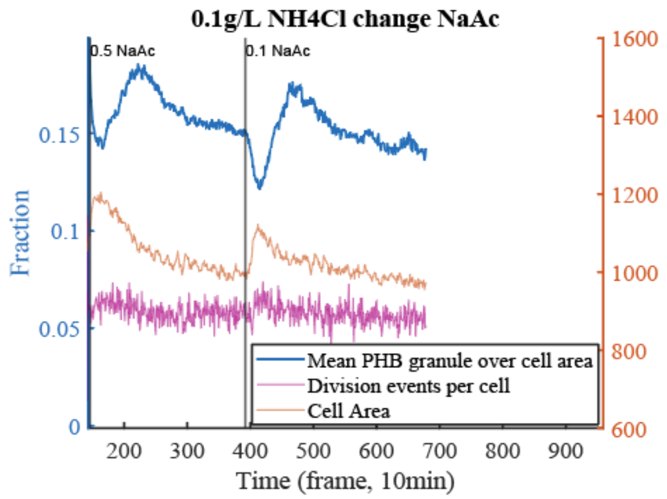


**S3. Integral feedback with additive vs multiplicative error**

Perfect adaptation, the return of a regulated variable to its baseline despite sustained changes in input, is a hallmark of integral feedback control. In this framework, the population-average PHB fraction, x(t) (granule area / cell area), is driven toward a target x* even when external nutrient levels u(t) (e.g., $NH_4Cl$) change abruptly. The essential feature is an internal controller state z(t) that integrates the error between the observed and target levels:

$$\dot{x} = s(u, z) - c(g), \qquad \dot{z} = -k_I \, e(x).$$

Here, s(u,z) is a slow synthesis or mobilization term, c(g) represents dilution and consumption coupled to growth g, and e(x) is the error signal. Two natural choices for error are:

- **Additive error**: $e_{\text{add}}(x) = x - x^*.$
  The system responds to the absolute deviation from the setpoint.

- **Multiplicative (fractional) error**: $e_{\text{mult}}(x) = \ln(x/x^*) \approx (x - x^*)/x^*.$
  The system responds to relative deviations, effectively sensing fold-change.

Both formulations enforce perfe[48,49]aptation (x→ x*) after step inputs[48,49]. However, they predict different transient behaviors. Additive error typically produces symmetric relaxations around the setpoint (fig.S3 b), while multiplicative error introduces asymmetry: equal fold-changes in input yield unequal additive deviations of x (fig.S3 c). In practice, when coupled with actuator saturation or differing time constants for synthesis and consumption, these models reproduce the main features of our data: return to baseline, dip–overshoot transients for nutrient step-ups, elevation–decay for step-downs, muted overshoot upon repeated perturbations (refractory), and diminished responses near the edge of growth permissibility.

This minimal formulation is meant as a phenomenological scaffold, not a molecular mechanism. It highlights how a simple integral controller can enforce robust setpoint regulation and how multiplicative sensing may underlie the asymmetries we observe. Multiplicative (fractional) error is biologically plausible, as fold-change detection is well documented in bacteria[63]chemotaxis and other sensory circuits[63]. Framed this way, the PHB system of *C. necator* joins a broader class of adaptive biological processes that responds robustly to fluctuating environments.

Resource-allocation theory predicts that bacteria often accept transient inefficiencies [41] maintain robust long-term regulation[41]. Perfect adaptation of PHB may reflect this principle—ensuring stable storage levels, even if it delays immediate growth optimization.

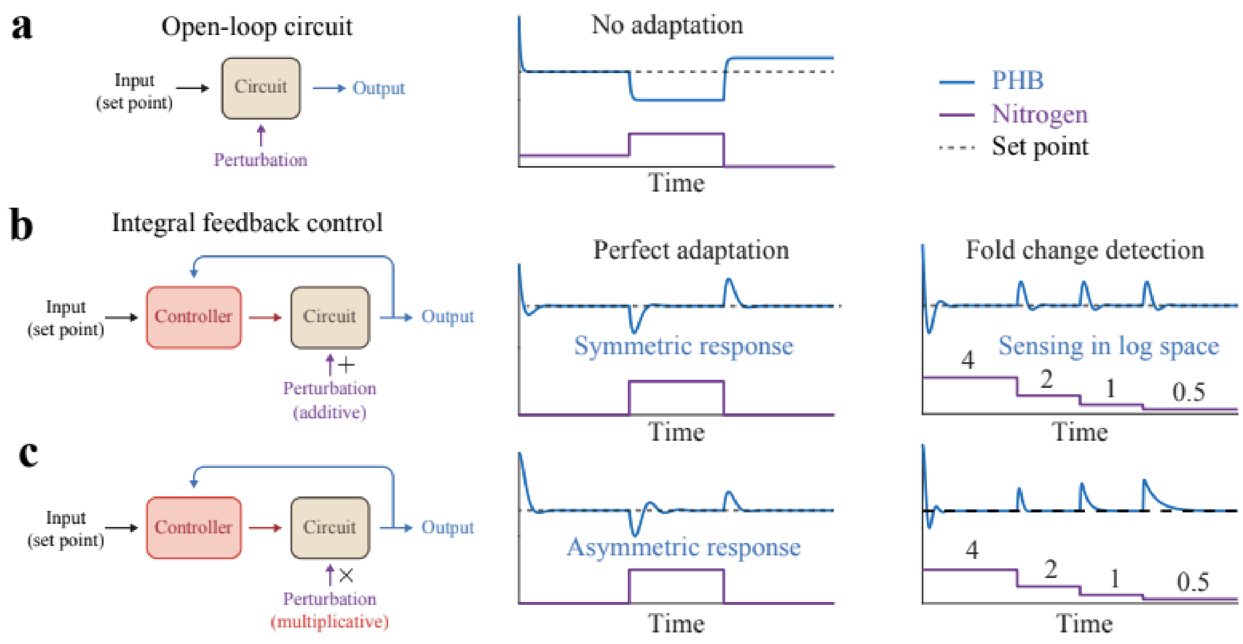


**S4. Full environment-by-species abundance heatmap and 2D hierarchical clustering.**

Heatmap showing the complete matrix of estimated abundance for PHB-associated microbial species across metagenomic environments queried with Branchwater (rows, environments; columns, species). Values represent the median estimated abundance within each environment, computed from genome coverage. Both axes were ordered by two-dimensional hierarchical clustering to highlight co-variation between environments and PHB-associated taxa. A reduced version of this analysis (subset of environments and species selected from the highest-ranked groups) is shown in Fig. 5d.

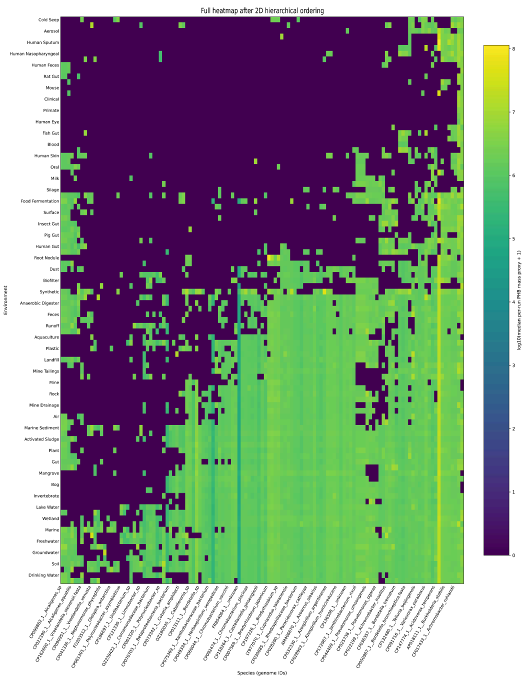


**Supplementary Fig. S5  Minimal segregation + growth model (single-lineage tracking; linear accumulation)**

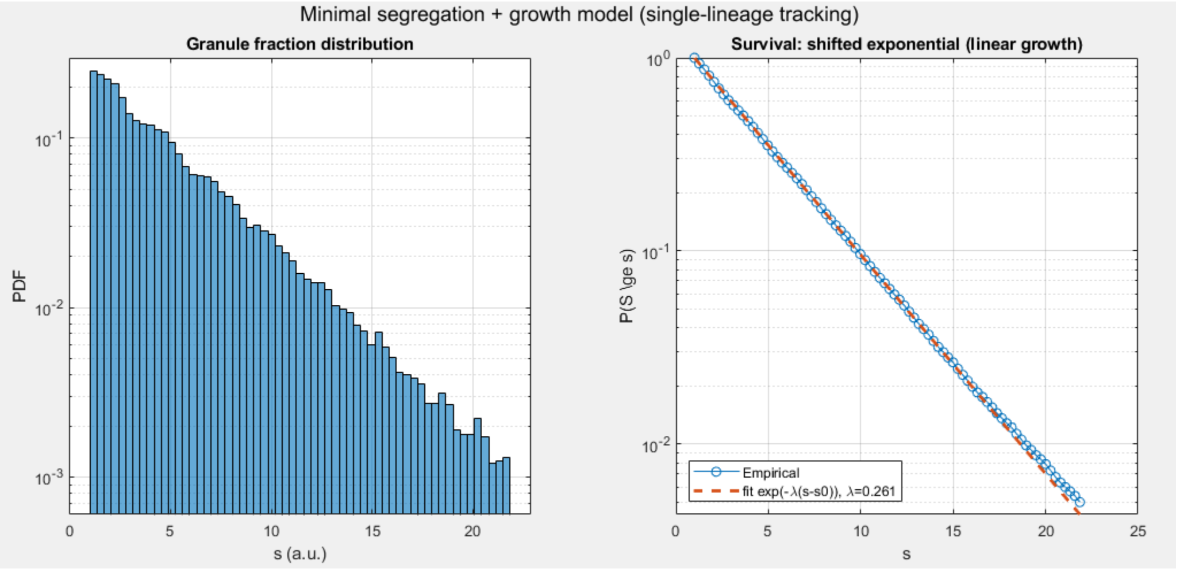


**a, Granule fraction distribution.**

Steady-state distribution of the granule-associated signal *s* obtained by sampling a tracked lineage at regular intervals after burn-in. The histogram is shown as a probability density (PDF) with a logarithmic y-axis to emphasize rare, large-*s* events. Upon granule acquisition, the signal is initialized at *s0* and then increases linearly with granule age; stochastic loss at division resets the lineage to a granule-free state and restarts the acquisition cycle.

**b, Survival analysis and shifted-exponential fit.**

Empirical survival function P(S >= s) for the same steady-state samples (semi-log plot), showing an approximately straight-line decay consistent with a shifted-exponential tail. The dashed curve is a maximum-likelihood fit of the form P(S >= s) ~ exp[-lambda*(s - s0)] for s >= s0, where *lambda* is estimated from the excess (s - s0).

**Supplementary Note S5 (model overview).**

The model describes a single tracked lineage that alternates between granule-free and granule-bearing phases. Granules are acquired after a stochastic delay Tacq (drawn per event), setting the baseline signal to s0. During the granule-bearing phase, the signal increases deterministically with granule age *a* as s = s0 + g*a. Cell divisions occur at stochastic intervals *tau*; at each division, the granule is retained in the tracked daughter with probability p_keep and otherwise is lost, terminating the bearing phase and initiating a new acquisition delay. This retention-at-division rule produces geometric persistence in generations and an approximately exponential persistence in time (set by p_keep and the mean division time), which under linear accumulation yields a shifted-exponential tail in *s* (panel b) and a long-tailed steady-state distribution (panel a).

**S6. Derivation of the analytic equal-fitness contour ln[b_CN(T_end)/b_PA(T_end)]=0.**

**Overview.**

Under periodic forcing with period P and duty cycle D, define the abundance duration per cycle as T_A = D*P. If the dynamics within each cycle depend only on T_A (and fixed strain parameters), then after n = T_end/P cycles the abundance ratio satisfies

$\ln[b_{CN}(T_{end})/b_{PA}(T_{end})] = n * \ln(M_{CN}/M_{PA})$,

where $M_{CN}$ and $M_{PA}$ are the per-cycle multiplication factors (i.e., the factor by which abundance changes over one complete cycle). The equal-fitness boundary $\ln[b_{CN}(T_{end})/b_{PA}(T_{end})]=0$ is therefore equivalent to $M_{CN} = M_{PA}$, and does not depend on $T_{end}$ except through the requirement that many cycles be simulated/considered.

Below we derive $M_{CN}$ and $M_{PA}$ for the nitrogen- and carbon-starvation regimes as implemented in the simulations, then solve $M_{CN} = M_{PA}$ for the threshold abundance duration DP0 (= $T_A$) and convert to a curve in the (D,P) plane via P = DP0/D.

1. **Nitrogen-starvation regime**

Implementation summary.
During abundance (R=1), CN grows at rate $r_{CN}$ continuously over the abundance window of length $T_A$ = D*P. PA grows at rate $r_{PA}$, but after each starvation-to-abundance transition it waits a fixed lag $\tau_{PA} = \ln(2)/r_{PA}$ during which it does not divide; after this lag, it resumes exponential growth at $r_{PA}$ for the remainder of the abundance interval. During starvation, neither strain grows.

Per-cycle multipliers.
CN grows exponentially for time $T_A$ each cycle, so

$$M_{CN} = \exp(r_{CN} * T_A).$$

PA grows only during the "post-lag" portion of abundance. The effective growth time is

$$G_{PA} = \max(0, T_A - \tau_{PA}),$$

so

$$M_{PA} = \exp(r_{PA} * G_{PA}) = \exp(r_{PA} * \max(0, T_A - \tau_{PA})).$$

Equal-fitness boundary.
Solve $M_{CN} = M_{PA}$:

$$\exp(r_{CN} T_A) = \exp(r_{PA} \max(0, T_A - \tau_{PA})).$$

This is a piecewise equation because of max(0, ...):

Case A: $T_A \le \tau_{PA}$. Then $\max(0, T_A - \tau_{PA})=0$, so $M_{PA}=1$ and the equality becomes $\exp(r_{CN}*T_A)=1$, which has only the trivial solution $T_A=0$. Thus, for any nonzero $T_A \le \tau_{PA}$, CN strictly outcompetes PA and there is no nontrivial equal-fitness boundary.

Case B: $T_A > \tau_{PA}$. Then $\max(0, T_A - \tau_{PA})=T_A - \tau_{PA}$, and the equality becomes

$r\_CN T\_A = r\_PA (T\_A - tau\_PA)$.

Rearranging,

$(r\_PA - r\_CN) T\_A = r\_PA tau\_PA$.

Substituting $tau\_PA = \ln(2)/r\_PA$ yields the closed form threshold

$DP0\_N = T\_{A,0} = \ln(2)/(r\_PA - r\_CN)$ (valid only when $r\_PA > r\_CN$ and $T\_{A,0} > tau\_PA$).

Therefore the analytic equal-fitness contour in the (D,P) plane is

$P = DP0\_N / D$.

(Here and below, D is restricted away from 0 because $P = DP0/D$ diverges as $D \rightarrow 0$.)

2. **Carbon-starvation regime**

Implementation summary.
At each abundance-to-starvation transition (R switches 1 -> 0), CN receives a one-time multiplicative fold-change of 1.5. At each starvation-to-abundance transition (R switches 0 -> 1), both strains enter a reduced-rate recovery phase at $r\_i/2$ for a fixed duration

$H\_i = t\_{rec,i} = 4*\ln(2)/r\_i$.

In the simulation implementation, the recovery timer is decremented continuously, so reduced-rate growth persists for exactly $H\_i$ even if the system enters starvation before the timer ends; after the timer ends, growth proceeds at full rate $r\_i$ only when R=1.

Per-cycle multipliers.
Let $T\_A = D*P$. Over a single cycle:

(i) During the recovery phase (duration $H\_i$), strain i grows at rate $r\_i/2$ irrespective of R(t), contributing a factor $\exp((r\_i/2) H\_i)$. With $H\_i = 4 \ln(2)/r\_i$, this simplifies to

$\exp((r\_i/2) H\_i) = \exp((r\_i/2)(4 \ln(2)/r\_i)) = \exp(2 \ln(2)) = 4$.

Thus, the reduced-rate phase contributes a fixed 4-fold multiplication per cycle for both strains (provided $H\_i$ completes within a cycle, which holds for the period grid used here, $P >= 10$ and $H\_CN \approx 10$).

(ii) After the recovery phase ends, strain i can additionally grow at full rate $r\_i$ for any remaining abundance time beyond $H\_i$, i.e. for $\max(0, T\_A - H\_i)$, contributing $\exp(r\_i*\max(0, T\_A - H\_i))$.

(iii) CN receives an additional multiplicative factor 1.5 at the abundance-to-starvation transition (one event per cycle for 0 < D < 1).

Therefore,

M_CN = 1.5 * 4 * exp(r_CN * max(0, T_A - H_CN)),
M_PA = 4 * exp(r_PA * max(0, T_A - H_PA)),

and the per-cycle ratio simplifies (the 4 cancels) to

M_CN/M_PA = 1.5 * exp( r_CNmax(0, T_A - H_CN) - r_PAmax(0, T_A - H_PA) ).

Equal-fitness boundary.
Set M_CN = M_PA, equivalently M_CN/M_PA = 1:

ln(1.5) + r_CNmax(0, T_A - H_CN) - r_PAmax(0, T_A - H_PA) = 0.

This is again piecewise due to max(0, ...). The relevant cases are:

Case A: T_A <= H_PA. Then both max terms are 0, so ln(1.5)=0 cannot be satisfied. Hence CN strictly outcompetes PA (no equal-fitness solution) in this regime.

Case B: H_PA < T_A <= H_CN. Then max(0, T_A - H_PA) = T_A - H_PA and max(0, T_A - H_CN)=0, so the condition becomes

ln(1.5) - r_PA*(T_A - H_PA) = 0,

giving

DP0_C = T_A,0 = H_PA + ln(1.5)/r_PA.

This solution is valid only if it lies in the assumed region, i.e. DP0_C <= H_CN.

Case C: T_A > H_CN. Then max(0, T_A - H_CN)=T_A - H_CN and max(0, T_A - H_PA)=T_A - H_PA, so the condition becomes

ln(1.5) + r_CN*(T_A - H_CN) - r_PA*(T_A - H_PA) = 0.

Using r_CNH_CN = r_PAH_PA = 4*ln(2), the H terms cancel and the expression reduces to

ln(1.5) - (r_PA - r_CN)*T_A = 0,

so

DP0_C = T_A,0 = ln(1.5)/(r_PA - r_CN).

This solution is valid only if it lies in the assumed region, i.e. DP0_C >= H_CN.

In practice, we compute DP0_C by evaluating the candidate thresholds in Cases B and C and selecting the one that satisfies its region condition; if neither condition is satisfied, no nontrivial equal-fitness boundary exists for that parameter set in the carbon-starvation model.

Finally, the analytic equal-fitness contour in the (D,P) plane is

P = DP0 / D,

with DP0 chosen as DP0_C (carbon-starvation) or DP0_N (nitrogen-starvation), and D restricted away from 0 (and away from 1 for carbon-starvation, because the 1.5 fold-change event is triggered only when an abundance-to-starvation transition occurs).

##### S7 genome completeness cutoff

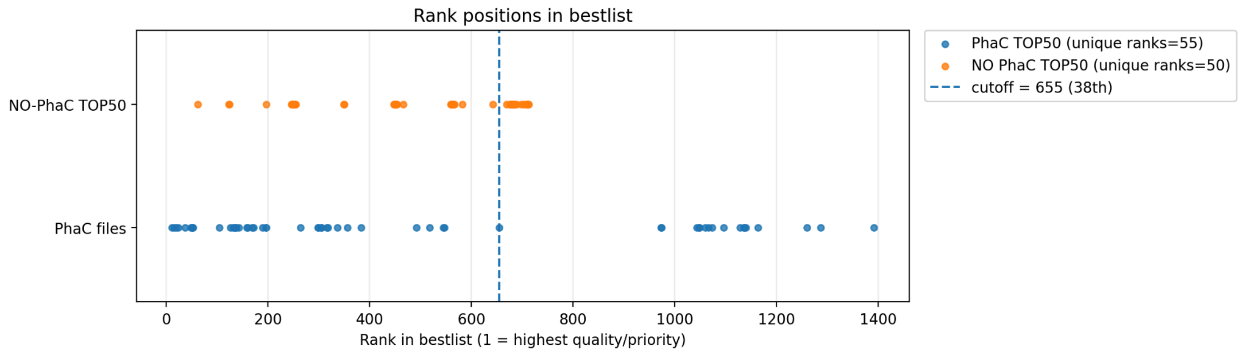


We identified 55 Betaproteobacteria species encoding PhaC and 50 species lacking PhaC, retrieved their genome assemblies from NCBI, and compared assembly quality across the two groups. To prioritize high-quality assemblies while maintaining consistent selection rules, candidate assemblies were ranked by (i) CheckM completeness in descending order, (ii) CheckM contamination in ascending order, (iii) assembly level in descending order (complete genome/chromosome/scaffold/contig mapped to 4/3/2/1), (iv) RefSeq status in descending order (RefSeq assemblies prioritized, with GCF_ preferred when available), and (v) contig N50 in descending order. We then applied an assembly-quality threshold chosen to yield matched sample sizes across groups: the threshold was set such that an equal number of species (n = 38) exceeded the threshold in each set (PhaC-positive and PhaC-negative), thereby controlling for assembly quality in downstream comparisons.
